# Comparing Machine Learning Approaches for Predicting CFD-Derived Stroke Risk Indicators in Atrial Fibrillation Patients

**DOI:** 10.64898/2026.06.04.730070

**Authors:** Paolo Melidoro, Riccardo Cavarra, Samia Mostafa, Gregory Y.H. Lip, Magdalena Klis, Steven E. Williams, Oleg Aslanidi, Adelaide De Vecchi

**Affiliations:** School of Biomedical Engineering and Imaging Sciences, King’s College London, United Kingdom; Institute of Life Course and Medical Sciences, University of Liverpool, Liverpool, United Kingdom; Department of Clinical Medicine, Aalborg University, Aalborg, Denmark, and Department of Cardiology, Lipidology and Internal Medicine with Intensive Coronary Care Unit, Medical University of Bialystok, Bialystok, Poland; Centre for Cardiovascular Science, The University of Edinburgh, Edinburgh, United Kingdom

## Abstract

Non-valvular atrial fibrillation (AF) is associated with a five-fold increased risk of stroke, mainly due to impaired contractility of the left atrium (LA) leading to blood stasis and subsequent thrombus formation within the left atrial appendage (LAA). Current AF stroke risk stratification schemes, such as the CHA₂DS₂-VASc/ CHA₂DS₂-VA score, use comorbidities and do not capture mechanistic factors like blood flow dynamics and hypercoagulability. To address this, we developed a multiphase computational fluid dynamics (CFD) model of the LA, incorporating patient-specific geometries; modelling of the coagulation cascade; and non-Newtonian blood behaviour within the LAA.

Using 84 simulation cases generated via Latin Hypercube Sampling of physiological blood parameters and 21 patient-derived LA anatomies, we trained surrogate machine learning models, including Ridge regression, XGBoost, Gaussian Process Emulators (GPEs), and deep learning networks, to predict CFD outputs such as blood viscosity in and fibrin concentrations in the LAA. Deep learning achieved R² values up to 0.90, with the accuracy increasing when both physiological parameters and the raw CT image were included. Other models showed uneven performance with R^2^ values below 0.7, highlighting the role of nonlinearities between parameters.

The study presents a novel CFD model that captures the transition from blood stasis to clot formation, representing the full thrombotic continuum underlying stroke risk in AF, and a deep learning approach to enable efficient prediction of mechanistic outputs of clinical value for stroke risk stratification in AF patients.

**Author Summary:** Atrial fibrillation is a common heart rhythm disorder that greatly increases the risk of stroke. In many patients, blood can pool inside a small pouch of the heart called the left atrial appendage, where clots may form and later travel to the brain. Current clinical tools used to estimate stroke risk mainly rely on a patient’s medical history and do not directly assess the mechanistic processes that lead to clot formation.

In this study, we developed a computer model that simulates how blood flows and clots inside the heart using patient-specific heart anatomies derived from medical imaging.

Our model combines blood flow, blood biochemistry, and the changing physical properties of blood during clot formation. We then used machine learning methods to predict these complex simulation results more efficiently. Deep learning models performed best, particularly when both clinical parameters and heart imaging data were included.

Our work provides a new way to study the full process linking abnormal blood flow to clot formation in atrial fibrillation. In the future, this approach could support more personalised and mechanistic assessment of stroke risk and help guide treatment decisions.

## Introduction

Atrial fibrillation (AF) is the most common cardiac arrhythmia and disrupts the regular contractile function of the heart seen in individuals with normal sinus rhythm [1], [2]. This loss of effective contraction reduces the heart’s ability to pump blood efficiently, resulting in blood stasis within the atria [3]. The left atrial appendage (LAA), a small pouch-like structure extending from the left atrium (LA), is particularly susceptible to this stasis and is the source of approximately 90 percent of thrombi responsible for ischaemic strokes in patients with AF [4].

To reduce the risk of thrombus formation, anticoagulation therapy is prescribed for AF patients considered to be at high risk, based on clinical risk stratification tools, with the most widely used being the CHA₂DS₂-VASc/ CHA₂DS₂-VA score. [2], [5] [6]. These schemes rely solely on clinical comorbidities and do not take into account biomarkers obtained from imaging or blood tests, despite evidence from studies showing that certain blood-based and image-derived biomarkers are independent predictors of stroke in patients with AF [7], [8]. As current risk stratification schemes have low efficacy in predicting stroke in patients classified as low risk, it has been suggested that their predictive accuracy could be enhanced by incorporating factors that reflect underlying mechanistic conditions, such as blood flow velocity in the LAA and markers of hypercoagulability, including fibrinogen concentration [9], [10].

One approach to incorporating blood biomarkers into stroke risk assessment is the ABC score, which includes NT-proBNP, an established marker of heart failure, to improve predictive accuracy [9]. However, this score does not include blood biomarkers of hypercoagulability, such as fibrinogen and peak thrombin concentrations, both of which are known to be elevated in patients with AF who have suffered a stroke [11], [12].

Geometric features of the LAA have also been associated with an increased risk of stroke. This includes links between stroke and certain LAA shapes such as wind sock, chicken wing, broccoli and cactus, as well as specific parameters like the distance to the first bend in the LAA anatomy and the LAA orifice area [13], [14]. While some studies have found that the chicken wing morphology is associated with a reduced risk of stroke and the broccoli morphology with an increased risk, classifying the LAA geometry into these four shapes is subjective. Significant variation has been reported in how clinicians categorise the LAA, suggesting that specific geometric parameters, such as those mentioned above, could offer a more practical approach to incorporating LAA features into stroke risk stratification [15], [16].

In addition to anatomical effects, the blood flow in the main LA cavity, and consequently inside the LAA, depends on the flow profile of the pulmonary veins (PVs). Subsequently a decrease in PV velocity has been linked with an increased risk of thrombogenesis in the LAA [17], [18]. Given the importance of these factors in the mechanisms of thrombus formation underlying stroke risk, it is essential to identify the most significant factors for risk assessment [6], [9], [17], [19]. A sensitivity analysis of the geometrical parameters of the LA has previously been conducted by Garcia-Isla et al. to determine which features have the greatest influence on blood stasis in the LAA [20]. However, this study performed local sensitivity analyses that did not fully capture the interaction between parameters that drive non-linear and non-additive processes. The use of computational fluid dynamics (CFD) to assess flow metrics in the LA for improving stroke prediction in patients with AF has become a widely researched area.

This approach has even been combined with biophysical models of coagulation to integrate the underlying mechanisms of thrombogenesis into CFD simulations [21], [22], [23], [24]. The application of CFD models in healthcare is hindered, however, due to the practical considerations associated with running such models. This includes the time required to run simulations, the costly computational infrastructure they demand, and the expertise in CFD that is rarely available within the healthcare environment.

Surrogate models have thus been proposed to predict CFD outputs at a fraction of the computational expense, by training the surrogate models using CFD outputs, including in applications involving the LA [25]. Approaches to creating surrogate models include utilising Gaussian Process Emulators (GPEs) and deep learning approaches [21], [26], [27].

This study therefore utilises a multiphase patient-specific CFD coagulation model to train machine learning models, which can then be used to predict the original CFD models output based on inputs including blood parameters, the LAA geometry and the PV velocities.

## Results

### CFD Model

The CFD simulations of the LA produced notable results, showing that the outputs were influenced by different input parameters. This was evident from the fact that the highest levels of fibrin accumulation at each location within the LAA, as well as the highest average viscosity in the LAA, occurred in different patients. The highest values observed were: 0.0177 Pa·s for average LAA viscosity; and fibrin accumulations of 5.9 × 10⁻¹⁰ moles at the orifice, 1.15 × 10⁻⁹ moles at the neck, and 1.58 × 10⁻⁹ moles at the tip. These simulations are illustrated below in Figure 1.

**Figure 1:**
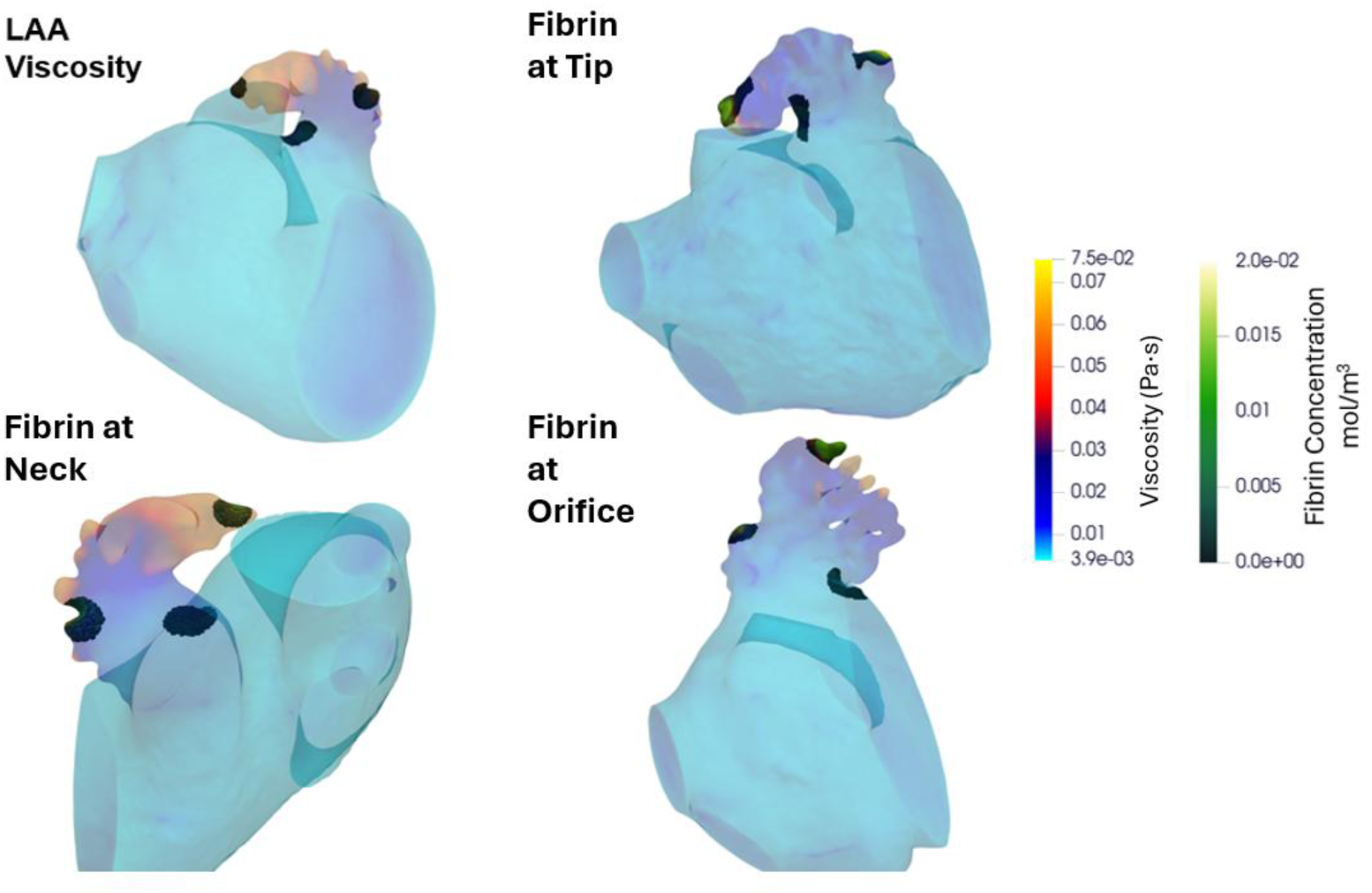
The simulation with the highest LAA viscosity is shown in the top left, while the other three simulations represent fibrin accumulation at each site where thrombin was initiated. The fibrin concentration at each site was summed spatially following six cardiac cycles to determine the total amount of fibrin present.

Figure 2 shows the temporal evolution of thrombin and fibrin in three regions of the LAA, measured at 0.2 s (early first cardiac cycle), 2.4 s (end of the third cycle), and 4.8 s (end of the sixth cycle).

**Figure 2:**
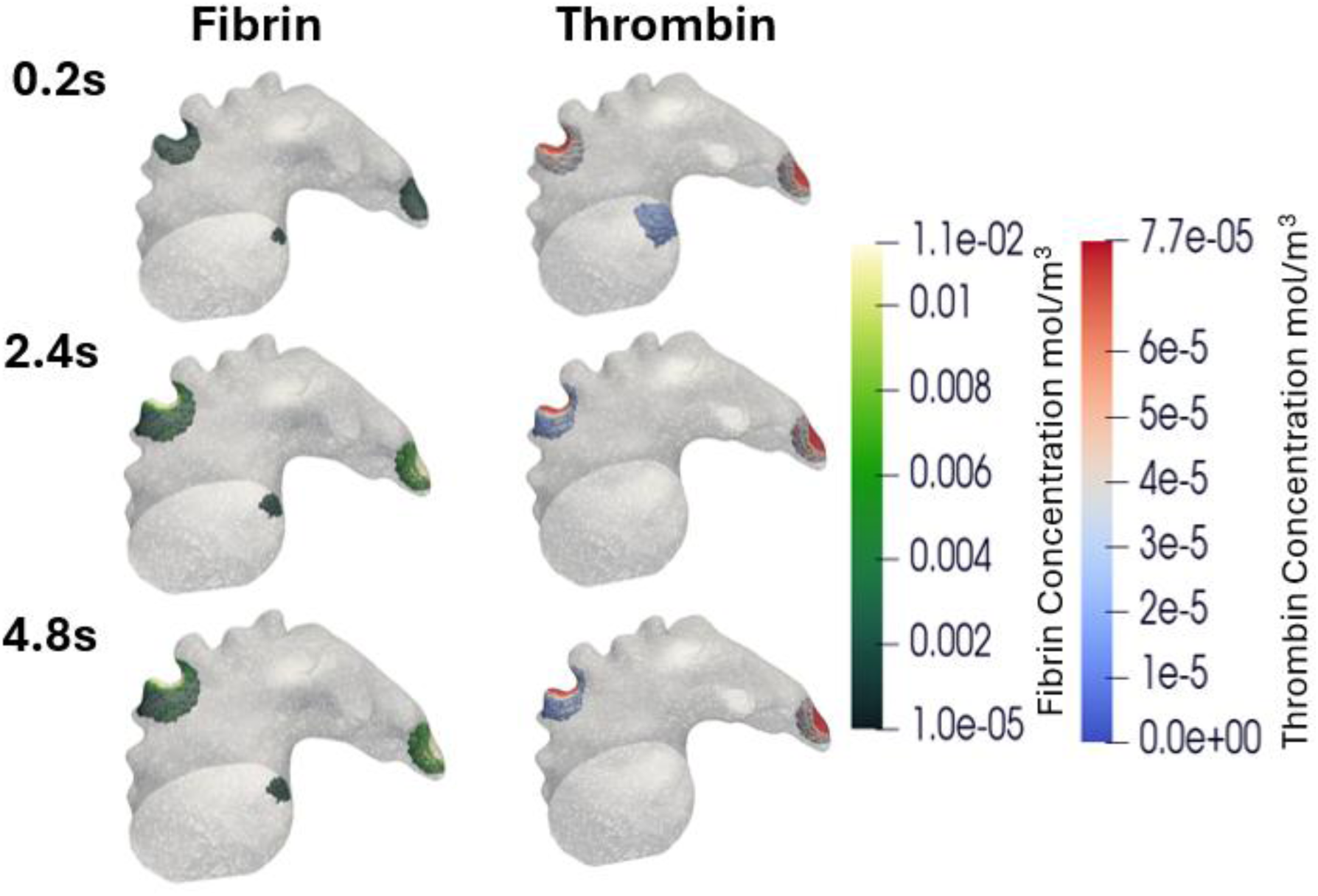
The temporal evolution of the fibrin and thrombin concentration within the LAA throughout the simulation, with the concentration of both shown at the start of the first cardiac cycle and then the end of the third and sixth cardiac cycle.

At the tip, thrombin increased gradually from 2.43 × 10⁻¹² mol at 0.2 s to 2.56 × 10⁻¹² mol at 4.8 s. Fibrin accumulated rapidly between 0.2 s and 2.4 s, rising from 5.13 × 10⁻¹¹ mol to 3.80 × 10⁻¹⁰ mol, and plateaued at 4.98 × 10⁻¹⁰ mol by 4.8 s. This indicates significant blood stasis at the tip, where flow is insufficient to prevent fibrin accumulation.

At the neck, thrombin decreased from 4.29 × 10⁻¹² mol at 0.2 s to 1.12 × 10⁻¹² mol at 4.8 s, while fibrin increased sharply from 9.12 × 10⁻¹¹ mol to 4.37 × 10⁻¹⁰ mol at 2.4 s before declining slightly to 3.54 × 10⁻¹⁰ mol at 4.8 s. The neck exhibited the largest spatial distribution of thrombin and fibrin, likely because flow was sufficient to convect thrombin and fibrin around the LAA but was insufficient to fully remove them. Although fibrin is distributed over a larger area at the neck, the tip accumulates the greatest total fibrin due to its higher local concentrations, which arise from the more extreme blood stasis in this region.

At the orifice, thrombin and fibrin remained low, decreasing from 8.05 × 10⁻¹³ mol and 1.52 × 10⁻¹¹ mol at 0.2 s to 1.79 × 10⁻¹⁶ mol and 5.9 × 10⁻¹² mol at 4.8 s, indicating that flow at the orifice is sufficient to prevent significant accumulation of fibrin and thrombin.

Figure 3 shows how the viscosity of the blood phase varies over the course of a cardiac cycle in the LAA. In this example LAA viscosity during the third cardiac cycle is visualised at 2.4 s, 2.8 s, and 3.2 s: blood viscosity exhibits a clear shear-dependent behaviour, with lower viscosity in high-velocity regions and vice-versa.

**Figure 3:**
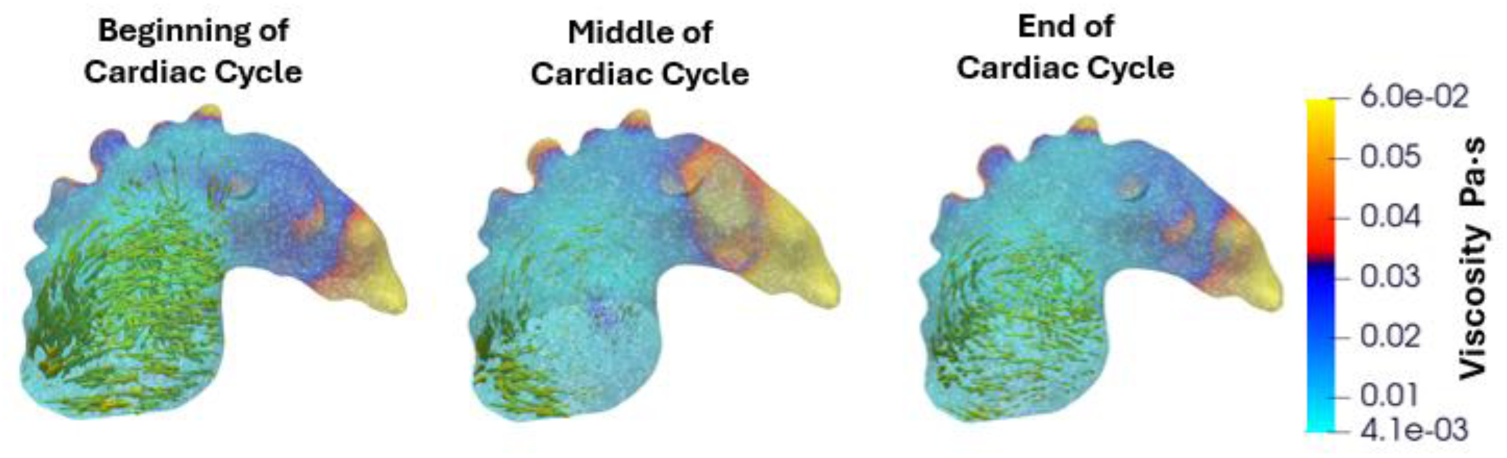
Temporal variation of the blood phase of viscosity within the LAA during the third cardiac cycle. The LAA viscosity is shown at the beginning (2.4 s), mid-cycle (2.8 s), and end (3.2 s) of the fourth cardiac cycle. Yellow arrows indicate velocity vectors, scaled according to local velocity magnitude.

At 2.4 s, viscosity is 0.012 Pa·s, reflecting the initial phase when flow into the LAA is elevated due to a delay between peak flow in the LA, which occurred in the middle of the previous cycle, and its propagation into the appendage. LAA viscosity peaks mid-cycle (2.8 s, 0.015 Pa·s), as the LAA flow at this point has declined, meaning there is a greater proportion of the LAA where the shear rate is low enough for non-Newtonian behaviour to occur. By 3.2 s, viscosity decreases slightly to 0.0116 Pa·s as the velocity in the LAA begins to increase again. It is also visible in Figure 3 that the highest viscosity consistently occurs at the LAA tip and within trabeculated regions, where shear is lowest.

### Multivariable Analysis

The multivariable analysis, as shown in Table 1, demonstrated region-specific associations between haematological and anatomical parameters and fibrin deposition within the LAA. At the orifice, fibrin deposition was positively associated with platelet count (β = 0.2423, p = 0.028), fibrinogen (β = 0.3950, p < 0.001), and haematocrit (β = 0.2365, p = 0.036), while pulmonary vein (PV) velocity showed a significant negative association (β = −0.3427, p = 0.002). At the neck, platelet count (β = 0.3253, p = 0.002) and fibrinogen (β = 0.4669, p < 0.001) remained positively associated, with PV velocity again inversely related (β = −0.3051, p = 0.004); other variables were not significant. At the tip, fibrin deposition was positively associated with platelet count (β = 0.2689, p = 0.013), fibrinogen (β = 0.4952, p < 0.001), and distance to the first LAA bend (β = 0.2575, p = 0.018), and negatively associated with LAA orifice size (β = −0.2566, p = 0.015), while PV velocity and haematocrit were not significant.

**Table 1:** Multivariable regression analysis of factors associated with fibrin deposition at different regions of the LAA and with average LAA viscosity.

| Input Parameter | Coefficient | P-Value |
| --- | --- | --- |
| Fibrin at orifice |  |  |
| Platelet count | 0.2423 | 0.028 |
| Fibrinogen | 0.3950 | 0.000 |
| Haematocrit | 0.2365 | 0.036 |
| PV Velocity | -0.3427 | 0.002 |
| Distance to LAA first bend | 0.0857 | 0.434 |
| LAA orifice size | -0.0291 | 0.783 |
| Fibrin at neck |  |  |
| Platelet count | 0.3253 | 0.002 |
| Fibrinogen | 0.4669 | 0.000 |
| Haematocrit | 0.0246 | 0.815 |
| PV Velocity | -0.3051 | 0.004 |
| Distance to LAA first bend | -0.0767 | 0.460 |
| LAA orifice size | -0.0298 | 0.766 |
| Fibrin at tip |  |  |
| Platelet count | 0.2689 | 0.013 |
| Fibrinogen | 0.4952 | 0.000 |
| Haematocrit | 0.1239 | 0.252 |
| PV Velocity | -0.1114 | 0.286 |
| Distance to LAA first bend | 0.2575 | 0.018 |
| LAA orifice size | -0.2566 | 0.015 |
| Average LAA Viscosity |  |  |
| Platelet count | -0.0967 | 0.384 |
| Fibrinogen | 0.3249 | 0.004 |
| Haematocrit | 0.3732 | 0.002 |
| PV Velocity | -0.3844 | 0.001 |
| Distance to LAA first bend | 0.0485 | 0.665 |
| LAA orifice size | -0.0952 | 0.381 |

For average LAA viscosity, fibrinogen (β = 0.3249, p = 0.004) and haematocrit (β = 0.3732, p = 0.002) were significant positive predictors, whereas PV velocity showed a significant negative association (β = −0.3844, p = 0.001); platelet count and distance to the first bend were not significant. Overall, fibrinogen showed the strongest and most consistent positive association with fibrin deposition across all regions, while PV velocity was consistently inversely associated, particularly at the orifice and neck, and anatomical factors demonstrated effects primarily at the LAA tip.

### Machine Learning Results

The predictive performance of all models across the four outputs from the CFD simulations is summarised in Figure 4, with the corresponding Pearson correlation scores shown in Figure 5. Overall, the models showed modest predictive accuracy, with none achieving an R² score above 0.7. Gradient Boosting performed best for Neck, achieving an R² value of 0.602, and the GPE also achieved a high R² of 0.576 for this output. GPE had the highest score for LAA Viscosity with R²=0.376, Ridge showed the highest R² for Orifice at 0.292, and Gradient Boosting led for Tip with R²=0.310, highlighting that all models still performed relatively poorly for predicting these outputs.

**Figure 4:**
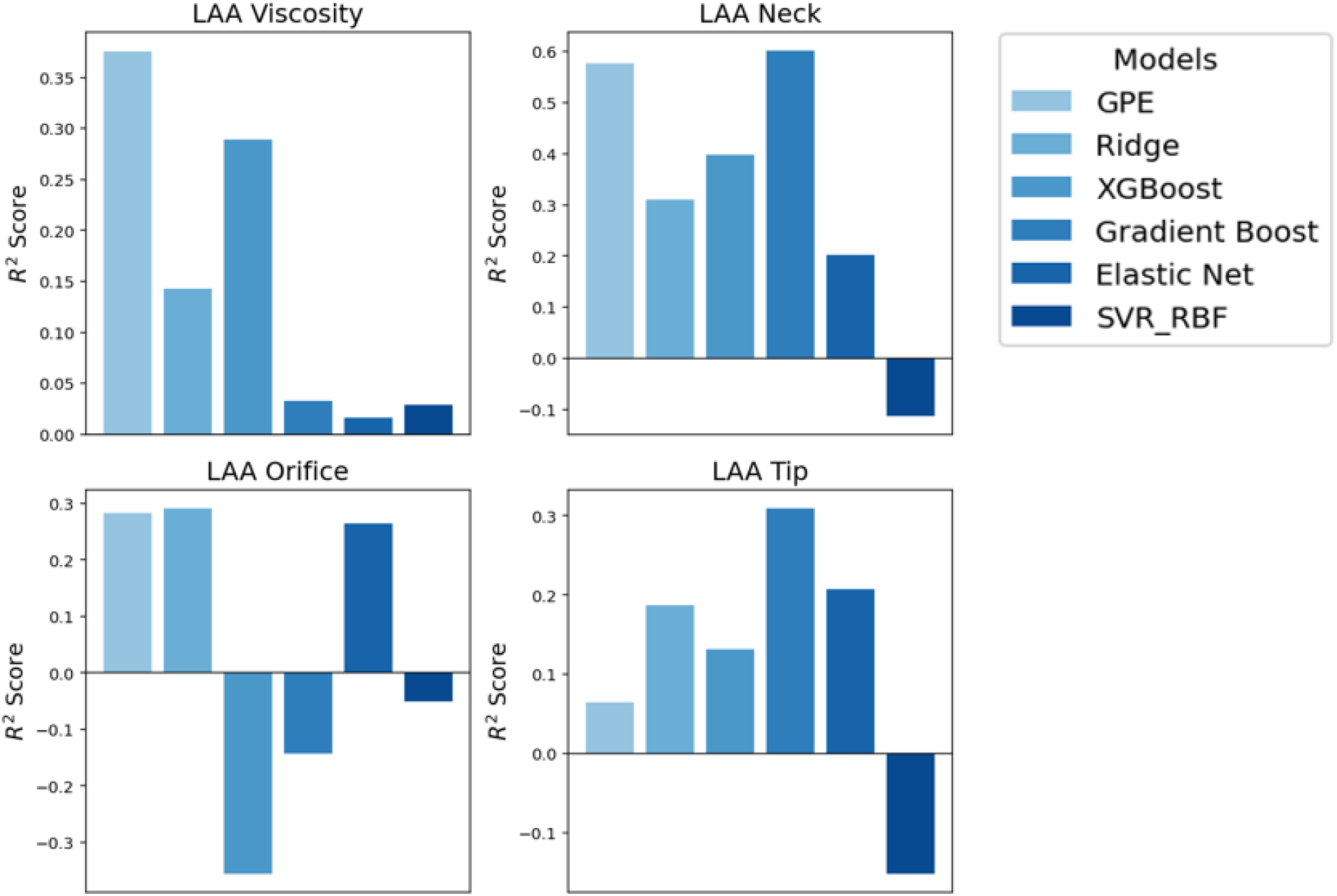
R² score for GPE, Ridge, XGBoost, Gradient Boosting, Elastic Net, and SVR_RBF models accuracy at predicting the CFD simulations output of average LAA viscosity and total fibrin at the neck, orifice and tip of the LAA.

**Figure 5:**
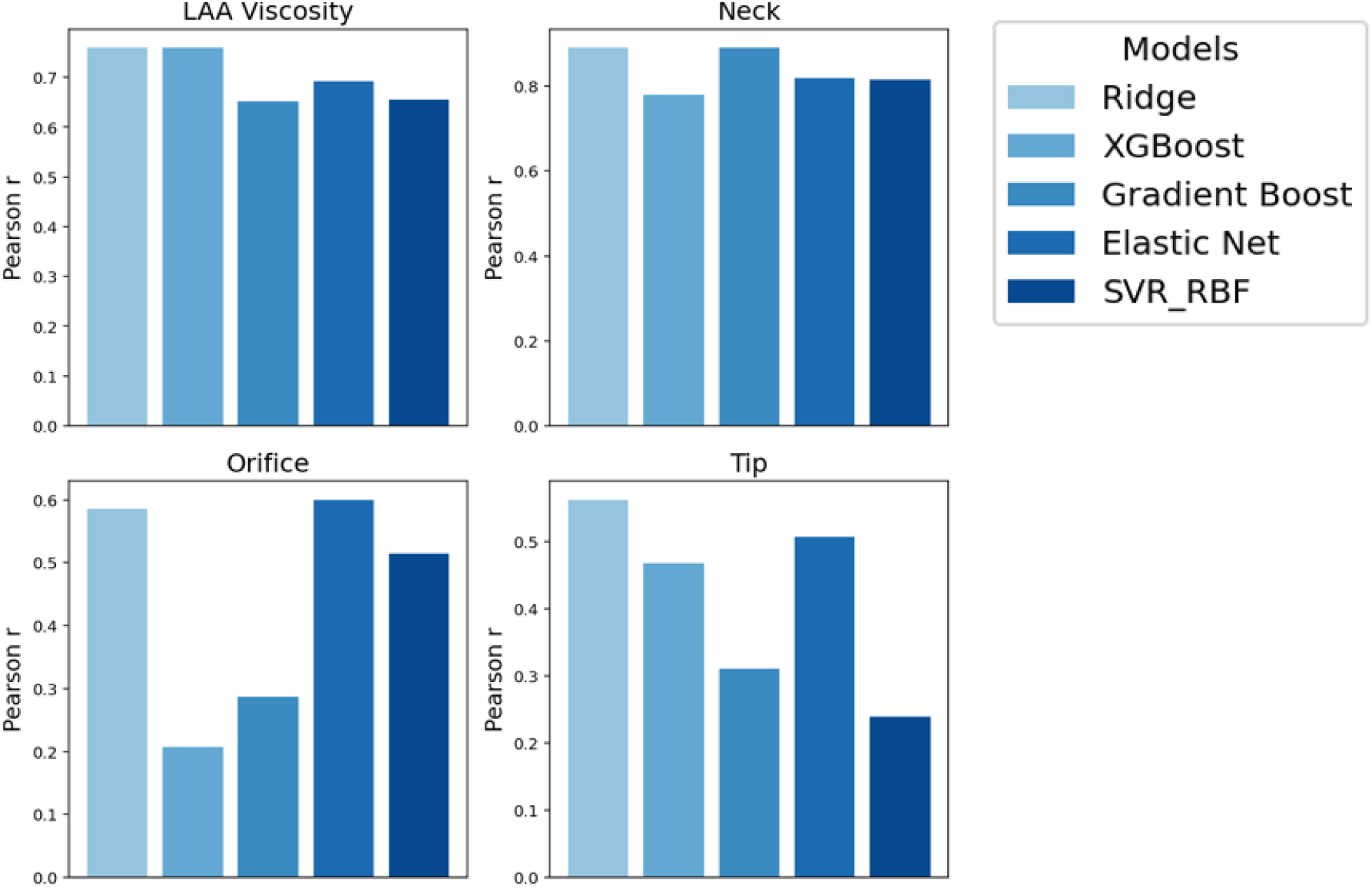
Pearson correlation coefficient for Ridge, XGBoost, Gradient Boosting, Elastic Net, and SVR_RBF models, quantifying the linear relationship between values predicted by the machine learning models and the actual CFD-simulated values for average LAA viscosity and total fibrin at the neck, orifice, and tip of the LAA.

Examining the Pearson correlations, which were calculated for each model except the GPE, Neck showed the strongest relationships, with Gradient Boosting achieving 0.890. LAA Viscosity also had relatively high correlations, led by XGBoost at 0.759. Orifice and Tip showed moderate correlations, with the highest values at 0.600 for Elastic Net (Orifice) and 0.562 for Gradient Boosting (Tip). Overall Gradient Boosting had the strongest correlations across the four outputs.

The limited R² values suggest that additional data or more features may be necessary to improve predictive accuracy and better reproduce the magnitude of the CFD outputs, though the models can identify the main relationships.

### Deep Learning Results

The predictive performance of different input combinations across the four CFD output parameters can be seen below in Figure 6. Using both image and all the available input parameters (“Image and parameters”) consistently yielded the highest R² scores across all outputs, reaching up to approximately 0.90 for LAA Blood Viscosity, 0.8 for both LAA Tip and Neck, and 0.7 for LAA Orifice. Models that incorporated all physiological parameters without images also performed well, with R² values above 0.65 for all outputs.

**Figure 6:**
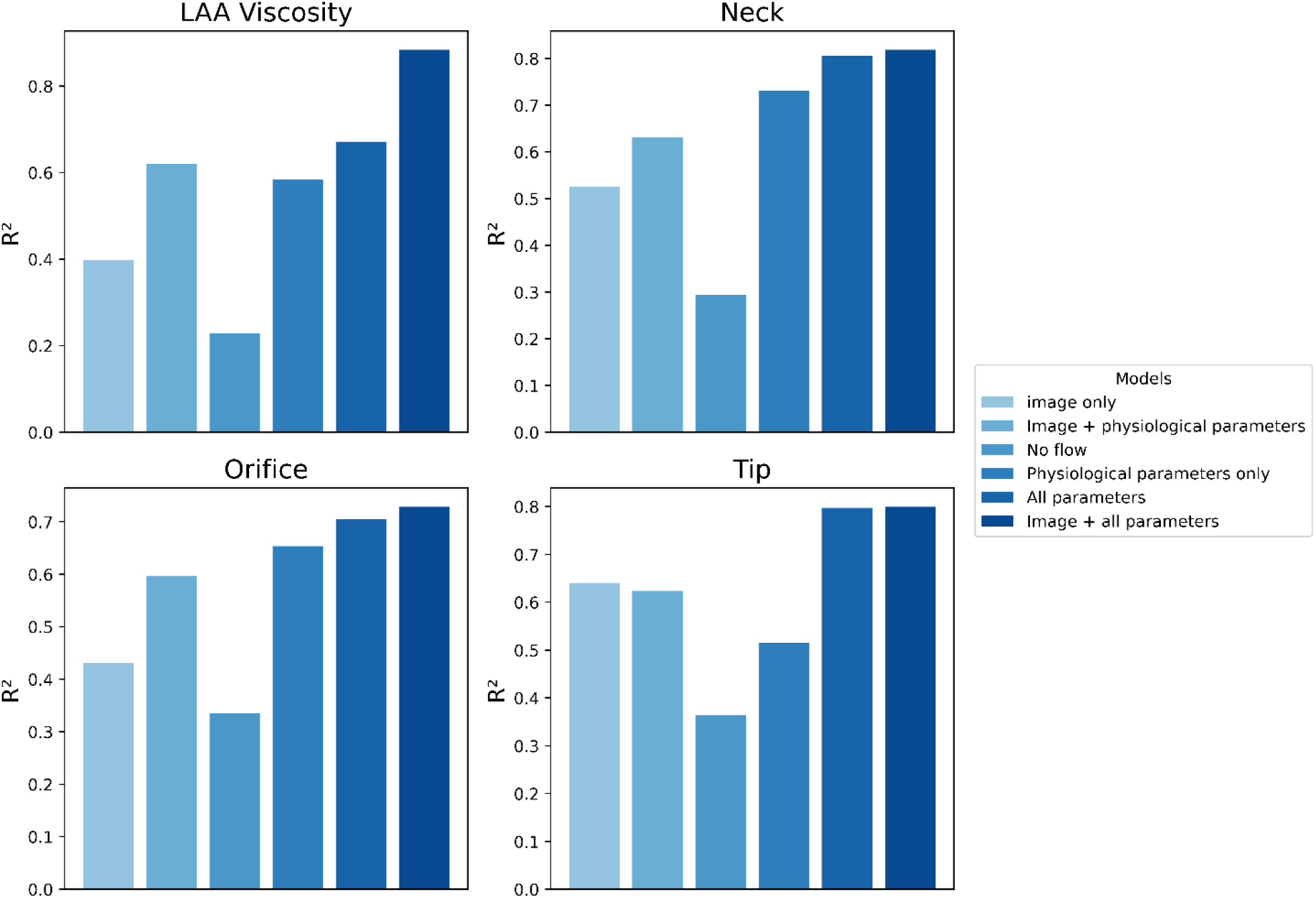
Predictive accuracy of the deep learning model for the four CFD outputs: LAA blood viscosity and total fibrin at the tip, neck, and orifice. The R² scores for each output are shown for different combinations of input parameters, as indicated in the legend.

In contrast, models using only images or only a selected few parameters alone showed noticeably lower predictive accuracy, with R² values ranging roughly from 0.30 to 0.65 depending on the output. The no flow combination of inputs showed the weakest performance for all outputs, indicating that combining these features without additional flow or parameter data may be insufficient.

These results outperform the other machine learning models, demonstrating the superior predictive accuracy of deep learning in this context. Notably, even when using only the same physiological parameters as the previous models, the deep learning approach achieves higher accuracy. This performance is further enhanced by integrating the raw CT images used to create the 3D LA into the model, highlighting the added value of combining imaging data with physiological parameters.

## Discussion

In this study, we present a novel implementation of a multiphase CFD model applied to patient-specific three-dimensional models of the LA. This approach enables the simultaneous representation of both the non-Newtonian behaviour of blood and the progressive increase in viscosity associated with fibrin accumulation during coagulation. It therefore provides a more comprehensive framework for modelling the full spectrum of the thrombotic milieu that develops during thrombogenesis within the LAA.

The non-Newtonian behaviour of blood is primarily caused by the fibrinogen mediated aggregation of red blood cells at low shear rates with LAA spontaneous echo contrast (SEC) also arising from this mechanism [28], [29], [30], [31]. LAA SEC is an imaging phenomenon observed using transoesophageal echocardiography and is a strong predictor of stroke risk in AF patients, with the increasing severity of LAA SEC correlating with a higher risk of stroke [32], [33].

LAA sludge represents a more advanced manifestation of SEC and is considered to confer a stroke risk comparable to that of established thrombus [34], [35]. It can therefore be interpreted as an intermediate stage between SEC, which is primarily driven by blood stasis, and fully developed thrombus, in which coagulation processes dominate [36], [37].

Our model captures this progression by demonstrating how viscosity initially increases due to stasis-induced non-Newtonian effects. Over time, viscosity continues to rise as a result of fibrin accumulation during the coagulation phase, ultimately leading to regions of elevated viscosity that may represent the transition towards clot formation.

Using multivariable analysis, we showed that different outputs from the CFD model were influenced by different inputs. At the orifice, fibrin is mainly associated with blood variables, particularly fibrinogen and platelet count, while the contribution from the LAA geometry was insignificant. The same is true at the neck, where blood effects dominate and anatomical variables, including distance to the first bend, show only weak and non-significant associations. In contrast, the tip is the only region where anatomy shows relevant associations, suggesting that geometry becomes more important further into the LAA. Fibrinogen was consistently a significant factor, while platelet count was significant in all cases except the LAA viscosity. PV velocity is inversely associated with fibrin deposition at the LAA orifice and neck but is not significantly associated at the tip. LAA viscosity is mainly associated with haematocrit and fibrinogen, as expected for non-Newtonian blood behaviour, and is inversely associated with PV velocity due to a lower PV velocity causing lower shear rates leading to stronger non-Newtonian, non-linear effects.

Even when the GPEs were tested using input parameter combinations that demonstrated the strongest correlations with the CFD-derived outputs, performance improved only moderately. This suggests that the emulator is unable to fully capture the complex nonlinear interactions governing fibrin deposition and viscosity within the LAA. For example, PV velocity likely interacts nonlinearly with fibrinogen concentration and haematocrit through the non-Newtonian rheological model that determines LAA viscosity. Such coupled relationships are difficult for the GPE to represent when assuming that each CFD output is a relatively smooth function of the supplied inputs.

This limitation became particularly apparent in the 5-fold cross-validation experiments, where simulation groups with the same anatomies were excluded from the training set.

By contrast, the deep learning model was able to learn more flexible nonlinear interactions between the same variables, resulting in consistently better predictive performance across folds.

This study also presents the first application of machine learning models, beyond GPEs, to predict coagulation-related outputs derived from CFD models of the LA, including XGBoost, deep learning, elastic net, and ridge regression models. The use of such models enables near-instantaneous prediction of stroke risk, while incorporating mechanistic information that is absent from conventional clinical risk scores such as CHA₂DS₂-VASc. This is particularly advantageous in healthcare settings where computational resources and specialist expertise required for CFD simulations are often limited.

The machine learning models trained in this study, excluding the deep learning approach, demonstrated limited predictive accuracy relative to the CFD outputs. This is likely attributable to the relatively small dataset, as reflected by the relatively high Pearson correlation coefficients. These findings suggest that while the models can capture general linear relationships, they struggle to accurately reproduce the magnitude of the outputs. Expansion of the dataset would therefore likely improve their performance.

In contrast, the deep learning model demonstrated more promising results, indicating that this approach can effectively predict CFD-derived outputs even with limited training data. This makes it particularly suitable for clinical contexts where large, high-quality datasets are difficult to obtain. Furthermore, the deep learning framework offers greater flexibility in integrating heterogeneous data sources. In this study, it enabled the incorporation of raw CT images through convolutional neural network layers for feature extraction. This addition improved predictive accuracy for three of the four outputs, further highlighting the advantages of a deep learning-based approach. However, despite these strengths, deep learning models are often limited by reduced interpretability, as their decision-making processes can be difficult to explain in clinically meaningful terms [38]. In addition, concerns regarding generalisability remain important, since models trained on relatively small or homogeneous datasets may not perform consistently across different patient populations, imaging protocols, or clinical centres.

An important consideration, however, in terms of practicality is which parameters would be necessary for the machine modelling as inputs to still achieve a high accuracy, and which imaging modalities would be required to derive these parameters. As seen in the results for the deep learning, the geometric parameters of the LAA, the PV velocity, and the use of raw imaging improve accuracy for most outputs. Each of these parameters, however, could be difficult to obtain in the context of stroke risk stratification in AF patients.

In this setting, investigations are typically limited to routine, non-invasive tests such as ECGs and transthoracic echocardiography (TTE). While ECG is universally performed, it provides no structural or flow-related information. TTE can assess LA size and cardiac function, but it has limited ability to accurately capture detailed LAA geometry [39]. In addition, although Doppler echocardiography can be used to measure PV velocity, obtaining accurate and reproducible measurements is technically challenging and highly operator dependent [40].

Cardiac CT can provide detailed anatomical information, particularly for LAA geometry, but its use is limited in routine stroke risk stratification due to its impracticality. CT imaging has inherent problems in this context such as cost and concerns around radiation exposure, especially given recent evidence linking widespread CT use with an increased population-level cancer risk [41]. Therefore, although these parameters improve model accuracy, their limited availability highlights an often-overlooked issue regarding the clinical practicality of utilising such imaging, particularly CT scans for stroke risk stratification in AF patients.

A potential solution is to use CT scans to generate the three-dimensional models for CFD simulations and then apply a machine learning approach to predict the outputs.

Instead of incorporating the raw CT image into the deep learning network, a TTE image from the same patient could be used to predict the output, allowing a clinically practical imaging modality to generate mechanistically informed predictions. However, this approach would still need to be validated against actual clinical outcomes, such as future stroke events, and compared with models that directly use TTE scans to predict stroke risk. Such a comparison would determine whether the mechanistic CFD simulation provides added value or is redundant. Notably, incorporating TTE data into patient models has already been shown to improve predictive accuracy compared to current stratification schemes [42].

A limitation of this study was the use of rigid-wall simulations. Numerous studies have shown that incorporating moving walls significantly affects flow metrics, with rigid walls tending to overestimate the severity of blood stasis within the left atrium [43], [44].

However, rigid-wall simulations can still provide useful information by representing a worst-case scenario, reflecting conditions when atrial contractility is at its lowest. Using dynamic imaging, such as cine CT, to create moving-wall simulations also has limitations. These images capture only a moment in time, and atrial contractility can vary significantly within the same AF patient. Consequently, the image may not represent the patient’s most impaired contractile state, and it is possible that the patient is in sinus rhythm when the image is acquired, meaning the contractility will not be impaired compared to when the patient is in AF [1].

Importantly, the goal of these models is not to detect thrombus at the moment the imaging is performed, but to predict whether a thrombus may develop in the future. In this context, a worst-case scenario simulation may actually be more informative. Future models could also integrate ECG data to estimate the most probable state of LA contractility by quantifying the degree of impairment in electrical activity, providing a patient-specific, dynamic prediction of stroke risk in AF patients.

Another limitation that has been previously mentioned is the limited size of the dataset; while using deep learning to some extent overcame this limitation future studies should use a larger dataset to examine whether the other machine learning model’s are able to achieve a high predictive accuracy when they have a complete dataset.

This study presents the first multiphase model of coagulation in the left atrium, capturing the full continuum of the thrombotic milieu, from LA SEC to fully formed thrombus in the LAA of AF patients, which underlies their elevated stroke risk. By providing a comprehensive mechanistic representation of thrombogenesis, this model offers a framework that could be used for stroke risk stratification in AF patients.

To make these simulations clinically feasible, various machine learning methods were employed to predict CFD outputs, drastically reducing computational expense. Among these, deep learning achieved the highest accuracy, with three of the four CFD outputs further improved when incorporating the raw CT images used to generate the patient-specific three-dimensional LA models.

Overall, this work represents an advance in both modelling and clinical translation. In terms of modelling, it combines coagulation processes with the non-Newtonian behaviour of blood in the LAA. Clinically, it demonstrates a practical approach to using mechanistic CFD-derived outputs for stroke risk prediction in AF patients. The application of deep learning to predict stroke-related CFD outputs provides a novel and practical approach that could be used in future to integrate mechanistic modelling of blood flow and thrombogenesis into current stroke risk stratification schemes, even creating virtual twin models and simulations [45].

## Materials and methods

The workflow of this study was as follows. First, 21 three-dimensional models of the left atrium (LA) were generated using contrast-enhanced CT scans of patients with AF, which had been previously segmented manually by a clinician [46]. Second, a new CFD coagulation model was used in each anatomy to generate synthetic datasets using Latin Hypercube Sampling (LHS) for key physiological parameters including PV velocity, fibrinogen concentration, haematocrit, and platelet count. The datasets were constructed so that each anatomy corresponded to 4 different LHS data points, resulting in a total of 21 x 4=84 simulation cases, of which 68 cases were used for training and 16 for testing. CFD simulations for each case incorporated both scalar transport and the biochemical reactions of the coagulation cascade [10], [47]. Finally, selected simulation outputs (viscosity and fibrin concentrations inside the LAA) were used to train a range of surrogate models, which were then tested to assess their accuracy.

### 2.1 Left Atrium Models

A total of 21 contrast-enhanced CT datasets from patients with AF were analysed, each acquired with a slice thickness of 0.5 mm. Detailed acquisition protocols for the CT scans are available from the SCOT-HEART investigators [46]. The images were segmented using Seg3D (NIH Centre for Integrative Biomedical Computing) to generate patient-specific geometries. These models were then post-processed, with surface smoothing carried out in MeshLab (add manufacturer), followed by further refinement in ParaView (add manufacturer), where the PVs and MV were clipped to isolate the region of interest.

### 2.2 CFD Model

To model flow in the LA, the commercial finite volume software ANSYS Fluent was used. A simplified sinusoidal function was applied as a boundary condition at each of the PVs, while the mitral valve was set as an outflow boundary condition, with all gradients normal to the valve set to zero. This simplification allowed the peak velocity to be easily scaled according to the Latin LHS values for this parameter, while preserving the pulsatile nature of PV flow. The peak PV velocity in the sinusoidal function was selected to reproduce the physiological range of velocities observed during the E-wave phase of PV flow in patients with AF (see Table 1).

Equal mass flow rates were prescribed at each of the four PV inlets to provide a simplified and controlled representation of LA inflow conditions, while the velocity magnitude within each vein was determined by its respective cross-sectional area. The range of applied flow conditions was based on physiologically realistic values reported for AF patients in the clinical literature and selected to capture inter-patient variability in LA haemodynamics.

Reaction-diffusion-convection (RDC) equations were used to focus on three primary biochemical species involved in the coagulation cascade: fibrinogen, thrombin and fibrin. These species were treated as scalar transport quantities, with the biochemical reactions incorporated through Fluent’s user-defined function feature [47], [48]. The RDC equations of the three biochemical factors and the reaction term for thrombin are as follows:

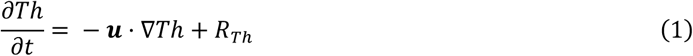

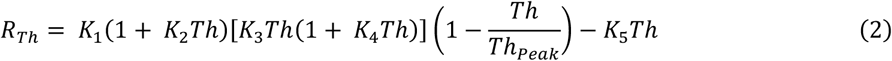

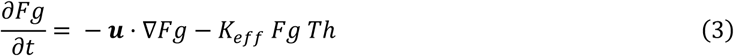

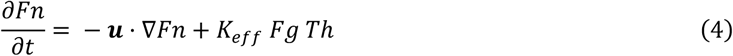

Where Fg, Th, and Fn are the concentrations of fibrinogen, thrombin and fibrin respectively, ***u*** is the flow velocity, *K_eff_* and *K_1-5_* the reaction rate constants as detailed in [22]. To initiate coagulation, thrombin was set to approximately 64% of its peak value to avoid modelling the entire thrombogram dynamics, which occur over tens of minutes. Three locations were selected as initiation sites for each anatomy: the tip, the neck and the orifice. These sites cover the vast majority of reported LAA clot locations, particularly those at the LAA tip [49]. Blood was modelled as a non-Newtonian behaviour fluid using a previously developed Carreau model of blood, dependent on fibrinogen and haematocrit [21].

To capture the dynamic changes in clot viscosity resulting from varying fibrin concentrations, a multiphase Volume of Fluid (VOF) approach was employed. In this framework, two distinct phases were defined: one representing blood and the other representing the clot. Each simulation was run for six cardiac cycles of flow time and took approximately 24 hours to complete on the 640 core SGI Alix-UV high performance computing cluster with Nehalem-EX architecture at King’s College London. The four outputs measured at the end of each simulation were the total fibrin accumulation at the three thrombin initiation sites, the orifice, the neck and the tip.

### 2.3 Gaussian Process Emulators

The points generated by the LHS fell within the physiological range for each parameter, as shown in Table 2. For fibrinogen, PV velocity and haematocrit, the inputs were directly produced by the LHS. However, for peak thrombin, the LHS generated values for platelet count, a more commonly available biomarker, which were then used to extrapolate peak thrombin levels based on experimental data [50].

**Table 2:** Ranges used for LHS values.

| Input Parameter | Range of values | Reference |
| --- | --- | --- |
| Fibrinogen | 5.88-17.6 $\mu\text{mol/L}$ | [12] |
| Haematocrit | 0.36-0.54 | [51] |
| Platelet Count | $150 \times 10^9$- $400 \times 10^9 \text{ L}^{-1}$ | [52] |
| Peak PV Velocity | 0.6-1.3 $\text{ms}^{-1}$ | [53] |

Each of the four outputs from the 68 simulations in the training dataset was used to train a separate GPE for each output using the GPErks software, and the models were then tested on the outputs of the 16 simulations from the testing dataset. [54]. All the GPE’s hyperparameters were jointly optimised by minimising the negative log-marginal likelihood using the GPErks emulation tool, which is based on the GPyTorch Python library and employs the ADAM optimiser [25], [55], [56]. Three different kernels, the Matern, Radial Basis Function (RBF), and Rational Quadratic (RQ) kernels, were tested for accuracy. Also included in the training of the GPEs were two geometric parameters of the LAA, the LAA orifice area and the distance from the LAA orifice to the first bend of the LAA, which were measured from the 3D anatomies of the LA using Spaceclaim (ANSYS).

Multivariable regression analysis was performed to further assess the independent associations between haemodynamic, haematological, and anatomical parameters and model outputs. All variables were entered simultaneously into the regression models to account for potential confounding effects. Standardised regression coefficients (β) and corresponding p-values were reported to quantify the strength and direction of associations, with statistical significance defined as a two-sided p-value < 0.05. Analyses were conducted using standard Python statistical libraries.

### 2.2 Other Machine Learning Models

XGBoost, Ridge, Gradient Boosting, Elastic Net, and Support Vector Regression with a radial basis function kernel (SVR-RBF) were trained and tested using the same dataset as the GPE to evaluate whether alternative machine learning approaches could better predict the CFD model outputs. To ensure consistency and comparability across all machine learning approaches, the input features were standardised using z-score normalisation prior to model training. Model performance was evaluated using 5-fold cross-validation, with the R² score used to compare predictive accuracy across the four CFD outputs. Additionally, the Pearson correlation coefficient was calculated to assess how well each model captured the linear relationship between the predicted and true outputs.

### 2.3 Deep Learning

Deep learning was employed to predict the same CFD outputs from the given inputs used to train the GPEs. All machine learning and deep learning models were trained and validated using the same data splits used to train the GPEs to allow direct comparison between methodologies. In addition, the raw 3D images used to generate the LA models were incorporated into the deep learning model through a convolutional neural network (CNN). Image pre-processing was minimal and involved normalisation and rescaling to homogenise the images. The 3D images were normalised to zero mean and unit variance, while rescaling with padding was performed to ensure that all input images had the same size while maintaining their original proportions. Similarly, the CFD input parameters were also normalised to zero mean and unit variance.

The CNN was trained to predict one target at a time; therefore, multiple CNNs with similar architectures were designed to predict each target for each input combination. The model followed a fusion architecture, where the CNN used a series of 3D convolutional blocks to downscale the images while preserving relevant information. Each convolutional block consisted of a 3D convolution, batch normalisation and ReLU activation, followed by spatial downsampling. The final feature map was flattened into a fixed-length latent representation using global average pooling before concatenation with the non-image inputs. The compressed image representation was then concatenated with the other input parameters and fed to a series of fully connected layers to make a prediction.

Dropout was used for regularisation, and grid search was used for hyperparameter tuning, selecting the number of layers, kernel size, learning rate and dropout rate that maximised accuracy during the validation stage. Training was performed on an RTX 4090 GPU using AdamW optimisation with Smooth L1 loss. Early stopping was used to interrupt training when the model started overfitting. The deep learning model architecture is represented below in Figure 7.

**Figure 7:**
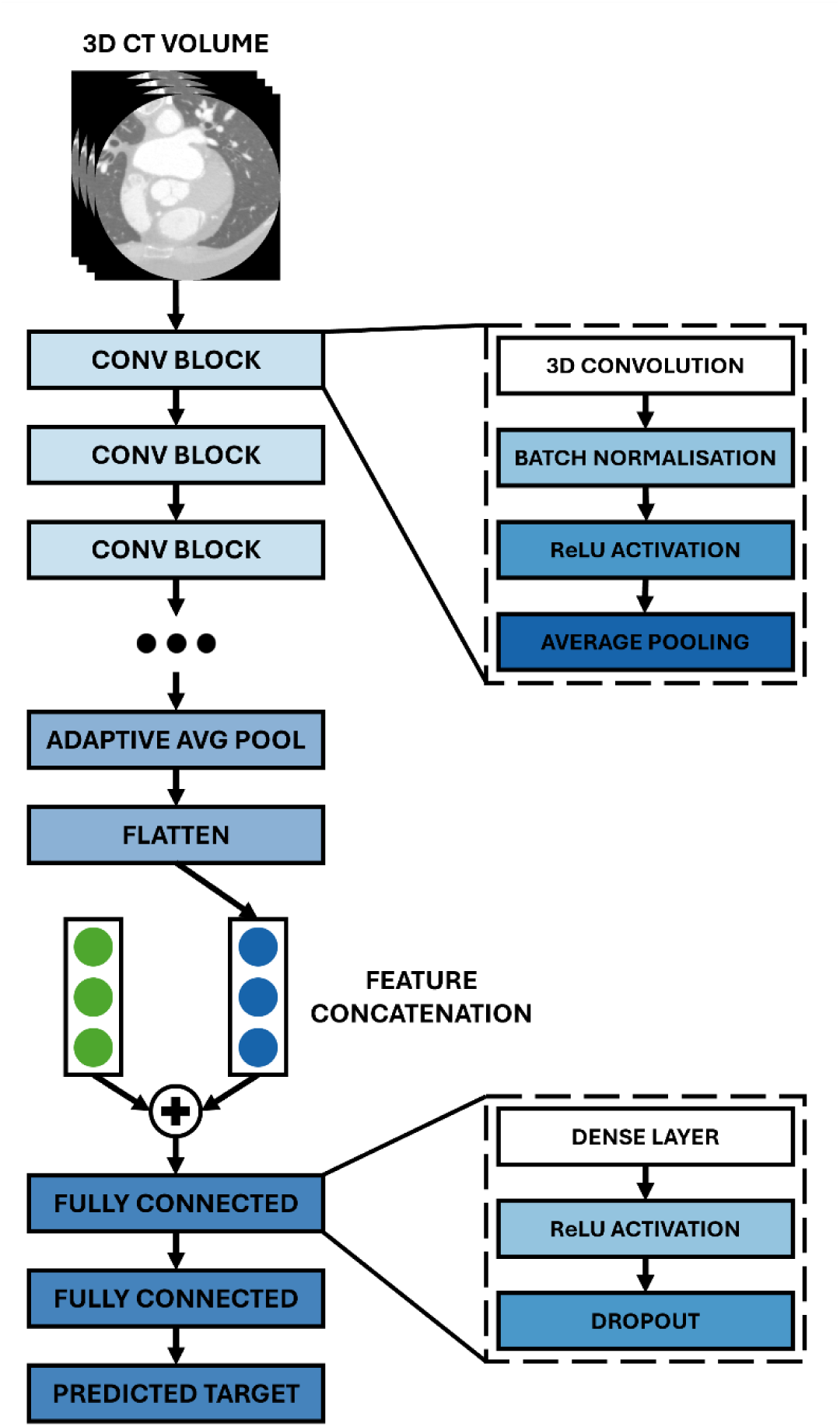
Deep learning model architecture. A series of convolutional blocks, comprising 3D convolution, batch normalisation, ReLU activation and spatial downsampling, were used to extract compact feature representations from the 3D CT images. The extracted imaging features were concatenated with the selected non-image input parameters, according to the specific training strategy, and passed through fully connected layers to perform regression. Hyperparameter tuning was used to optimise the architecture for each regression task, including the number of layers, kernel size, learning rate and dropout rate.

The accuracy of the deep learning model was quantified using the R² score and evaluated across different combinations of inputs, as shown in Table 3. Performance was compared against the corresponding machine learning models to assess whether incorporating raw 3D image information improved predictive accuracy.

**Table 3:**
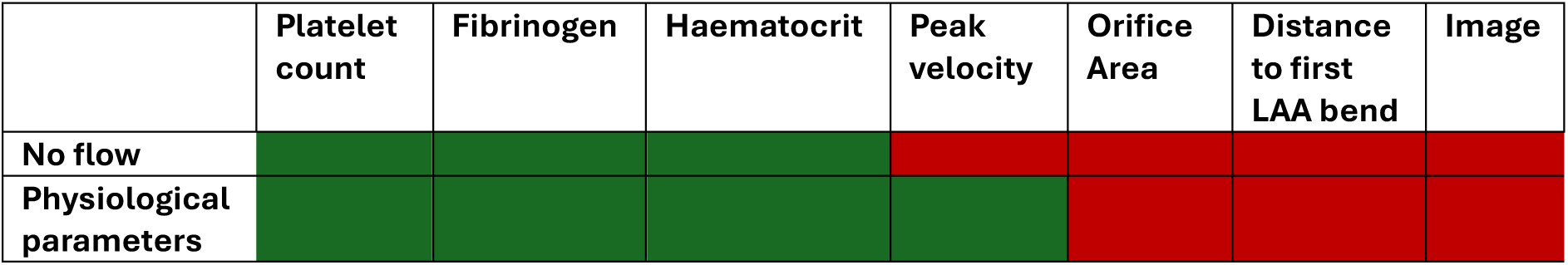
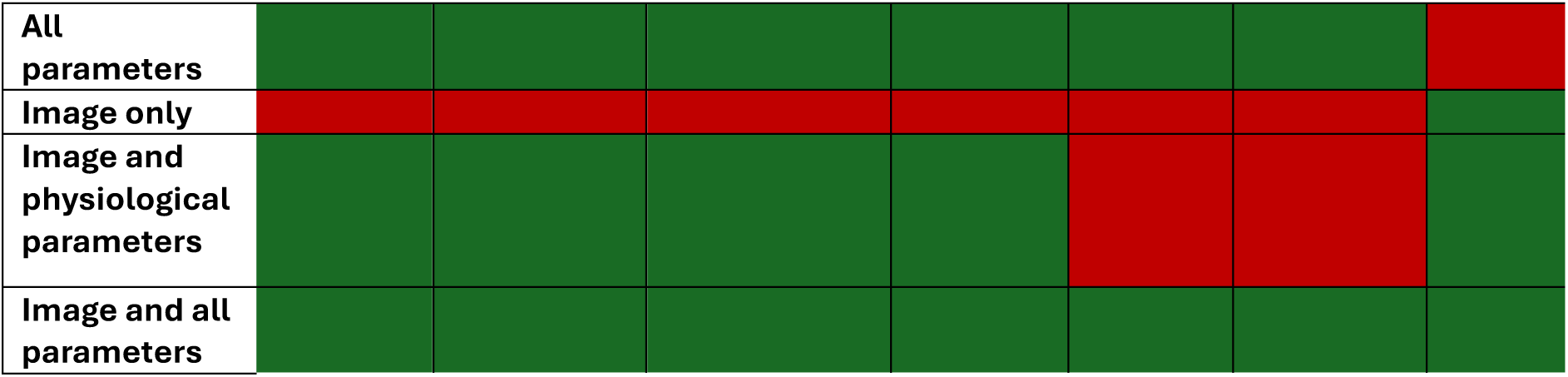
Combinations of inputs used for training the deep learning network, red indicated the input was excluded while green indicated the input was included. A total of six different combinations was used: no flow, physiological parameters, all parameters, image only, image and physiological parameters, and image and all parameters. This was done combining seven different inputs: platelet count, fibrinogen, haematocrit, peak velocity, orifice area, distance to first LAA bend and the raw CT image.

